# Hippocampal place cell sequences during a visual discrimination task: recapitulation of paths near the chosen reward site and independence from perirhinal activity

**DOI:** 10.1101/2024.04.18.590059

**Authors:** P. Marchesi, J. Bos, M. Vinck, C.M.A. Pennartz

**Affiliations:** Cognitive and Systems Neuroscience Group, SILS, University of Amsterdam, Netherlands; Research Priority Program Brain and Cognition, University of Amsterdam, Amsterdam, the Netherlands; Donders Institute for Brain, Cognition and Behavior, Radboud University and Radboud University Medical Centre, Nijmegen, The Netherlands; Ernst Strüngmann Institute for Neuroscience in Cooperation with Max Planck Society, Deutschordenstraße 46, 60528 Frankfurt, Germany

**Keywords:** Expectancy, Memory, Outcome, Parahippocampal cortex, Replay

## Abstract

Compressed hippocampal place-cell sequences have been associated with memory storage, retrieval and planning, but it remains unclear how they align with activity in the parahippocampal cortex. In a visuospatial discrimination task, we found a wide repertoire of hippocampal place cell sequences, which recapitulated paths across the task environment. Place cell sequences generated at reward sites predominantly reiterated trajectories near the chosen maze side, whereas trajectories associated with the side chosen in the previous trial were underrepresented. We hypothesized that neurons in the perirhinal cortex, which during the task display broad firing fields correlated with the animal’s location, might reactivate in concert with hippocampal sequences. However, we found no evidence of significant perirhinal engagement during virtual trajectories, indicating that these hippocampal memory-related operations can occur independently of the perirhinal cortex.

## Introduction

The hippocampus (HPC) is known for its causal role in the formation of episodic memories, but over time animals and humans come to rely less on this structure when applying memorized information to cognitive and behavioral decision-making (Eichenbaum, 2000; Moscovitch et al., 2005; Tse et al., 2007; Winocur et al., 2010). Although (Marr, 1971) famously postulated that the hippocampus transfers mnemonic information to the neocortex during sleep – an idea followed up by the two-stage model of memory consolidation (Buzsáki, 1989, 1996) – the mechanisms by which the HPC communicates with nearby medial temporal lobe structures and distant cortical sites remain poorly understood. One of the obstacles to gaining deeper insight into these processes is that we do not know how activity in parahippocampal structures, such as the perirhinal cortex (PER), is coordinated with hippocampal memory operations (Burwell and Amaral, 1998; Buzsáki, 1996; Doan et al., 2019; Eichenbaum, 2000; Fiorilli et al., 2021; Fiorilli et al., 2024; Miyashita, 2019; Ruikes et al., 2024; Suzuki and Amaral, 1994). It is becoming increasingly clear that the PER serves as an important hub in the medial temporal lobe, potentially controlling the formation of associative memories in the sensory neocortices via its projections to layer I (Doron et al., 2020). Alternatively, parahippocampal and other neocortical structures may process mnemonic information without necessarily interacting with the HPC (Kaefer et al., 2020; O’Neill et al., 2017).

During sleep or rest epochs, the HPC replays sequential neural representations of the animal’s trajectory following behavioral experience (Kudrimoti et al., 1999; Skaggs and McNaughton, 1996; Wilson and McNaughton, 1994). CA1 replay occurs prominently during population burst events (PBEs) associated with sharp-wave ripples, which originate in the HPC (Buzsáki, 2015; Suzuki and Smith, 1988), are causally relevant for spatial memory (Ego-Stengel and Wilson, 2010; Girardeau et al., 2009; Jadhav et al., 2012) and are thought to act as excitatory events driving replay in cortical and subcortical target regions of the HPC (Jadhav et al., 2016; Ji and Wilson, 2007; Pennartz et al., 2004; Rothschild et al., 2017; Rusu and Pennartz, 2020; Sirota et al., 2003; Wierzynski et al., 2009).

Hippocampal replay also occurs in the wakeful state, especially during periods of reduced mobility (Karlsson and Frank, 2009), which is of special interest because awake replay may reveal mechanisms subserving initial memory formation and memory retrieval to guide cognitive search, planning and decision-making during ongoing behavior (Carr et al., 2011; Diba and Buzsáki, 2007; Foster and Wilson, 2006; Pezzulo et al., 2014; Pfeiffer and Foster, 2013). During wakeful PBEs, hippocampal sequences can follow the direction of animal locomotion (‘forward replay’) or occur in temporally reverse order, which has been reported especially for reward sites (Foster and Wilson, 2006). In wakeful subjects, hippocampal ensembles can replay trajectories close or remote to the subject’s current position (Karlsson and Frank, 2009), express new sequences the animal has never experienced before (Gupta et al., 2010) and can be modulated by reward delivery (Ambrose et al., 2016; Bhattarai et al., 2020; Singer and Frank, 2009). Whereas reverse replay may enable retrospective evaluation and memory consolidation, especially in relation to reward delivery (Foster and Wilson, 2006; Kalenscher and Pennartz, 2008; Shin et al., 2019), forward replay may support sampling of future possibilities (Pezzulo et al., 2014; Pfeiffer and Foster, 2013; Shin et al., 2019; Wu et al., 2017). In addition to forward and reverse replay, which are thought to be occurring mostly during hippocampal sharp-wave ripples (SWRs), virtual trajectories generated by hippocampal neurons have also been reported in association with theta sequences, which are also compressed in time and can occur during locomotion as well as near-immobility, e.g. at decision points on a maze (Foster and Wilson, 2007; Johnson and Redish, 2007).

As yet, little is known about the coordination of virtual trajectories between the HPC and parahippocampal regions functioning as intermediate network nodes towards more distant neocortical structures. Although a previous study reported coordinated replay between area CA1 and the deep layers of medial entorhinal cortex (MEC; Ólafsdóttir et al., 2016), this replay was found during sleep. In contrast, O’Neill et al. (2017) found replay in MEC superficial layers occurring independently of the HPC, which raises the question whether other parahippocampal areas such as PER coordinate their replay activity with the HPC.

We analyzed neuronal ensemble activity simultaneously recorded from hippocampal area CA1 and PER to investigate whether virtual trajectories are manifested as a coordinated process between these two areas in a behavioral setting with distinct task phases. We identified trajectory events based on PBEs which have been associated with hippocampal SWR activity (Davidson et al., 2009; Wu et al., 2017) but may occasionally occur during theta sequences; to avoid confusion we will therefore mainly use the term ‘trajectory events’ instead of (classic, SWR-associated) ‘replay’. The behavioral task was a visual discrimination task set on a figure-8 maze which allowed us to establish the relationship between trajectory events and the display of visual images based on which decisions were made, arrival at reward sites and reward consumption, as well as to a resting state during intertrial intervals (ITIs).

We identified a wide variety of single-area CA1 trajectory events during the different phases of the task. First, we found no evidence for enhanced occurrence of place cell sequences when visual images were shown, whereas we did observe an increased occurrence during ITIs. Second, hippocampal sequences generated at reward sites generally reflected paths on the same side of the figure-8 maze that the rat was on, whereas paths traveled on the side chosen on the previous trial were generated significantly less often. Third, PER activity was correlated with the position of the animal during the task, such that it could be exploited to decode the arm of the maze currently occupied. Thus, we hypothesized that perirhinal spatial representations would be evoked during CA1 sequences in a coordinated recapitulation of recent experience. However, we found no evidence that hippocampal trajectory events significantly engaged PER activity in an inter-areal memory operation.

## Materials and Methods

### Behavioral task, recording technique and data acquisition

Rats were trained to perform a two-choice visual discrimination task on a figure-8 maze. Each trial began with an inter-trial interval (ITI) randomly lasting between 15-25 s, during which the animal was confined to the middle arm by way of two removable barriers (Fig. 1A, B). After a short sound cue, the rat could trigger the onset of visual stimuli by breaking an infrared photobeam. Two visual stimuli (CS+ and CS-) appeared on the two monitors located in front of the central arm of the maze (Fig. 1A; the side of appearance of each stimulus was determined by a pseudorandom sequence which guaranteed a balance between left and right trials). The transparent barrier (Fig. 1A) through which the rat could see the visual stimuli was removed 4.2 seconds after stimulus presentation, and the rat could choose one of the side arms. After the animal passed a point of no return, the arm was blocked such that the animal could no longer walk back to the opposite arm, and the decision was considered final (a correct decision corresponding to the side of the CS+). Upon a correct trial, rats received two or three pellets (BioServ, dustless precision pellets, 14 mg) at the reward site on the corresponding arm, plus an additional one after the rat returned to the middle lane. Rats performed significantly better than chance.

**Figure 1.**
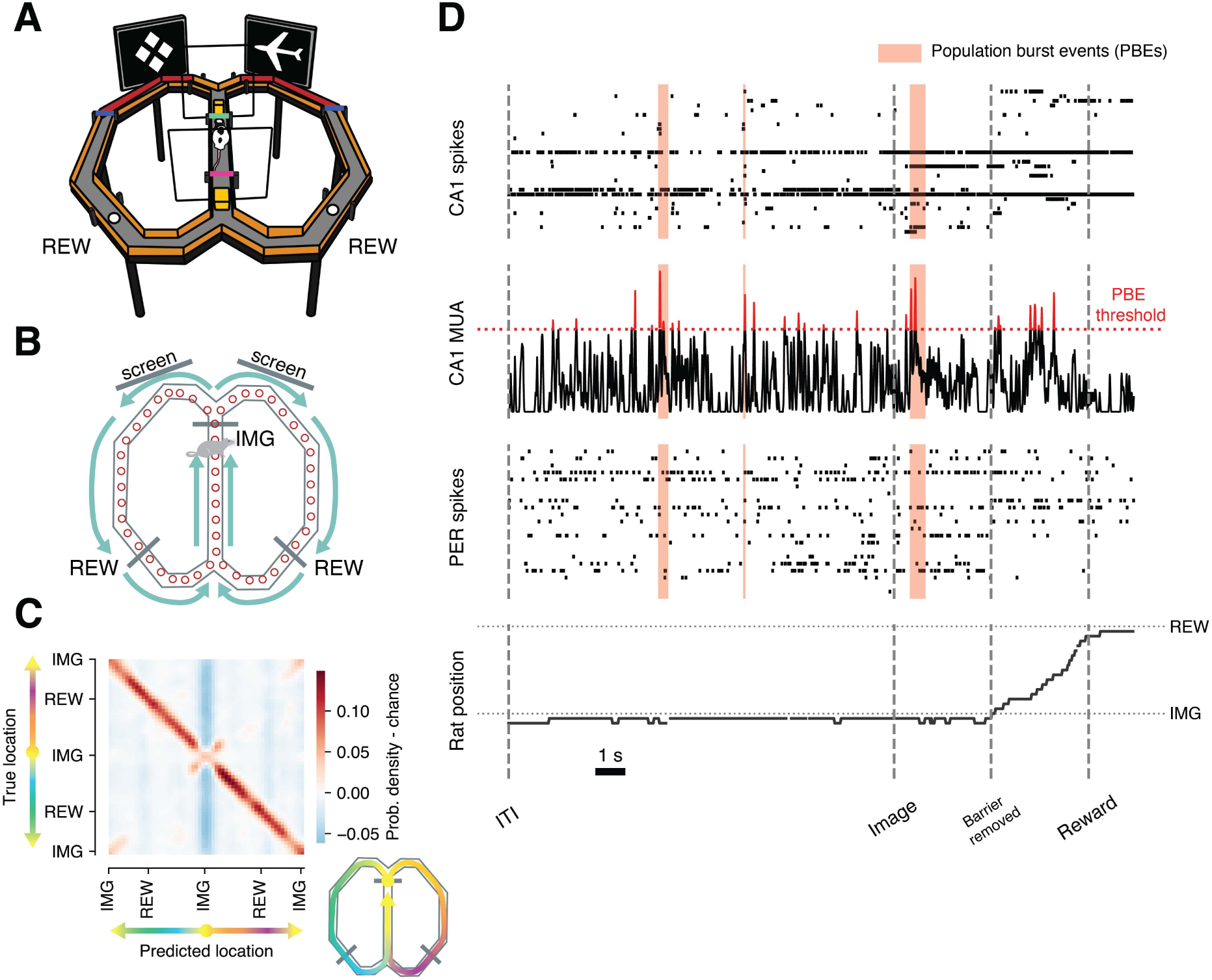
Simultaneously recorded ensembles in hippocampal area CA1 and perirhinal cortex. (A) Two-choice visual discrimination task on the figure-8 maze. REW: reward site. The green horizontal bar on the central arm indicates the front barrier (with transparent frame). The magenta horizontal bar behind the rat indicates the rear barrier. **(B)** Schematic view of the figure-8 maze from above, with red circles indicating the spatial bins used to discretize rat location. IMG: location of rat when facing visual stimuli. Arrows indicate locomotion direction during task performance. **(C)** Confusion matrix (averaged across sessions, N=16) for the decoding of actual animal location from CA1 ensembles. Darker red shades along the diagonal indicate higher probability that time bins in which the animal is at a given location are assigned by the decoder to the correct location, compared to a shuffled distribution. Decoding was significantly better than chance in all 16 sessions considered. **(D)** Rasters of simultaneously recorded ensembles in area CA1 and PER during the execution of a trial. Hippocampal multi-unit activity (CA1 MUA) was used to detect population burst events (PBEs, red shaded bands), which were subsequently analyzed to see if they contained place cell sequences. ‘Barrier removed’ indicates the moment at which the rat was allowed to start running along the arms of the maze from the decision point towards the reward sites and marks the end of the Image phase. The period from barrier removal until the arrival at reward sites was not analyzed due to consistently high locomotion speeds.

For the analysis of virtual trajectories we defined three task phases. The ‘ITI’ began once the animal had left the reward site in the previous trial and lasted until the visual stimuli for the upcoming trial appeared on the screens positioned in front of the maze. The ‘Image’ phase marked the time during which the animal was exposed to the visual stimuli, up until the removal of the front barrier (which allowed the animals to start running along one of the maze arms). Lastly, the ‘Reward’ phase consisted of the time after the animal had passed a photobeam placed at the reward site (and received the food pellets, in the case of correct trials), and before it left the reward site to return to the central arm. When the animals travelled from the front barrier to the reward sites, they were consistently running along the arms of the maze, generally without stopping until they reached the reward site, therefore these periods were excluded from further analysis.

Data was collected from three male Lister Hooded rats. Custom-built microdrives containing 36 individually movable tetrodes (quad-drives) were implanted into each rat’s right hemisphere (Lansink et al., 2009; Vinck et al., 2016). These included eight recording tetrodes directed to the PER (area 35/36; skull coordinates: -5.0mm anteroposterior (AP) and 5.0mm mediolateral (ML (Paxinos and Watson, 1998)), eight to the dorsal hippocampal CA1 area (-3.5mm AP and 2.4mm ML), eight to the somatosensory cortex (S1BF: -3.1mm AP and 5.1mm ML) and eight to the visual cortex (V1M, -6.0mm AP and 3.2mm ML to bregma, with one additional tetrode per area that could be used as a reference). Recordings performed in S1BF and V1 were not analyzed here. Neural activity was recorded with a 144-channel Digital Neuralynx Cheetah set-up (including 16 reference channels; Neuralynx, Bozeman MT) using tetrodes (Gray et al., 1995) (Nichrome wire, California Fine Wire, diameter: 13 mm, gold-plated to an impedance 500–800 kΩ at 1 kHz). LFPs were continuously sampled at 2,035 Hz and bandpass-filtered between 1 and 500 Hz. In order to identify single units, spikes were sorted based on waveform peak amplitude, energy, and first derivative of the energy using a semi-automated clustering algorithm followed by manual refinement (KlustaKwik, by K. Harris, and MClust 3.5, by A.D. Redish). Clusters with more than 0.1% of inter-spike intervals of less than 2ms were not accepted. The task setup was filmed at 25 Hz during recordings, and an array of light-emitting diodes placed on the headstage allowed for offline tracking of the animal’s location. The recordings we considered for this analysis yielded a total of 557 single units in CA1 and 238 in PER; further subselections were applied depending on the analysis performed. For further details on data acquisition and histology, see (Bos et al., 2017).

### Data preprocessing

#### Position data

Animal position was extracted from the video and cleaned from tracking artefacts using custom software. Speed was computed from the x and y position across video frames and smoothed with a Gaussian window with sigma equal to 4 frames (160 ms). The maze was subdivided in 51 spatial bins (Fig. 1B).

#### Sharp-wave ripple detection

LFP data was filtered between 105 and 300 Hz. Harmonics of residual line-noise was removed using a band stop filter (148-152 Hz). The absolute trace was smoothed with a Gaussian kernel (0.015 s duration, 0.003 s standard deviation.). Next, peaks were detected which exceeded four standard deviations (standard deviation of the non-smoothed data) for at least 6 ms. Durations of high frequency events around the peaks were determined by taking the times the signal surpassed a 1.33 std threshold. Peak events with overlapping durations were combined. Any high frequency periods shorter than 30 ms were excluded. If during any 2 s period more than 15% of the time was classified as high frequency event, all events in this period were excluded. Similarly, any high frequency period with signal peak exceeding a threshold of 2000 (uV, unsmoothed) was excluded as a noise peak. All remaining high frequency events were considered sharp-wave ripples (SWRs).

#### Classification of neuron types

Cell types were identified based on spike waveforms (Vinck et al., 2016). Waveforms were centered on the peak and normalized between 1 and -1. Plotting peak-to-trough duration and repolarization at 0.45 ms after the peak revealed three different sub-groups: fast spiking (presumably interneurons), broad spiking (presumably pyramidal cells) and unclassified cells.

### Decoding of rat position

#### Data selection

Decoding of the rat’s position (or the maze arm the rat was on) from CA1 and PER neurons was performed in all sessions containing at least 20 single units. This amounted to 16 sessions including CA1 and 11 sessions including PER. We used all cells which fired at least 30 spikes across the whole task phase. For hippocampal decoding we further selected only putative pyramidal neurons.

#### Data processing

Population vectors were constructed by binning spiking data in 400 ms bins. The position of the rat on the maze was linearly interpolated at the centers of the time bins, and assigned to the nearest bin (if the nearest bin was within 15 cm from the interpolated position, otherwise the sample was removed). Similarly, speed was linearly interpolated between the centers of the time bins. Time bins with no spikes, or during which locomotion speed of the animal was lower than 12 cm/s, were excluded from the decoding of actual animal position during task performance (Fig. 1C and Supplementary Fig. 1).

#### Bayesian decoding

We implemented a Bayesian decoder following (Davidson et al., 2009; Zhang et al., 1998), which determines the probability distribution of rat locations estimated from population vectors using Bayes’ rule:

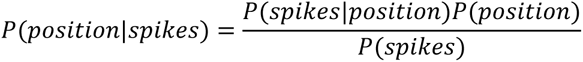

Assuming a uniform prior *P*_(*position*)_ and because the total probability across all positions must sum to 1, we only needed to compute the likelihood *P*(*spikes*|*position*) We first generated the rate maps *f_i_*(*x*) for every unit *i* and location *x*. The rate maps were smoothed using a Gaussian filter with a standard deviation of 1 bin. Then, for every time window of duration τ, we constructed the vector *K* = (*k*_1_, . . . , *k*_*N*_) containing the number of spikes fired by each unit, from 1 to N, during the interval. Assuming the firing rates of the neurons are independent from each other and follow Poisson statistics, the likelihoods *P*(*spikes*|*position*) are given by:

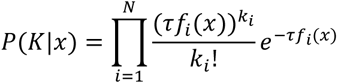

Thus

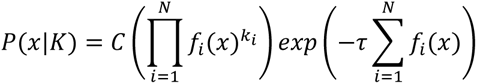

Where *C* is a normalizing constant which depends on τ and the number of spikes fired. The output of the decoder is a one-dimensional probability density function of location, which is independent of past (or future) estimates of locations (i.e. memoryless). The maximum-likelihood estimate of position was then taken as the predicted position.

#### Cross-validation

We employed a stratified k-fold cross-validation with 3 folds (i.e., the validation preserved the percentage of samples for each class in each fold) to split the data into training and test sets, and constructed the rate maps *f*_*i*_(*x*) using the training sets.

#### Evaluation of decoding performance and decoding density maps

For every CA1 recording, we gathered the predictions from all cross-validation folds and computed the decoding error as the average Euclidean distance between predictions and true locations. We then generated a distribution of surrogate errors by randomly permuting the predictions 500 times, and considered decoding to be significantly better than chance when the proportion of surrogate errors smaller than the observed error was below 0.0001. To visualize hippocampal decoding of position, for a given recording session we generated a confusion matrix with the cross-validated predictions, and normalized it over the rows (i.e. the true labels; Fig. 1C), such that every row represents the probability that the decoder assigns a firing rate vector recorded at a given location to all the other possible locations. We then generated 500 bootstrap-normalized confusion matrices by randomly permuting the predicted locations, and subtracted their average from the normalized confusion matrix. Lastly, we smoothed the debiased confusion matrix with a Gaussian filter (standard deviation of 1 spatial bin), and averaged across all sessions before plotting. The same procedure was used for CA1 and PER decoding of maze arm identity (Supplementary Fig. 1) where the Euclidean error was replaced by the balanced accuracy score, defined as the average of recall (ratio of true positives over all positive instances) obtained in each class, and no Gaussian smoothing was employed.

### Detection of virtual trajectories with Bayesian decoding

#### Data selection

We selected recordings which contained at least 20 CA1 neurons, for a total of 16 sessions. The trajectory analysis was performed using only putative pyramidal neurons which fired at least 15 spikes during the task. The decoders were trained on all time points with at least 1 spike and in which locomotion speed of the animal was higher than 12 cm/s.

#### Detection of Population Burst Events

The multi-unit activity (MUA) of putative pyramidal cells in hippocampal area CA1 was divided across 1 ms bins and smoothed with a Gaussian kernel with a standard deviation (s.d.) of 20 ms. We identified population burst events (PBEs) as periods where MUA exceeded 3 s.d. above the mean of the whole task period (Davidson et al., 2009). We defined the start and end of each period as the time points in which the MUA crossed the mean, and further adjusted its bounds to 2 ms before the first spike and 2 ms after the last spike within the event. We then identified a subset of PBEs as candidate trajectory events based on several criteria. We first discarded events for which the average movement speed of the animal was higher than 3 cm/s and which lasted less than 50 ms or more than 2000 ms. Events for which the locomotion speed could not be reliably determined were discarded. We further excluded events with fewer than 4 active pyramidal cells and in which less than 10% of the population was active. For every event, we determined the average X and Y position of the animal during the time of the PBE, and assigned it to the nearest spatial bin. Although PBEs showed significant overlap with SWRs (44%), we also found PBEs during non-SWR epochs; both SWR and non-SWR related PBEs were included here.

#### Decoding of Population Burst Events

We followed an approach similar to O’Neill (O’Neill et al., 2017), but because in our case the task imposed a specific running direction, we did not distinguish between forward and reverse trajectories by building a separate rate map for each running direction. Thus, we only fitted two Bayesian decoders, one for all left trials and one for all right trials. Spikes within each candidate trajectory event were binned with a time window of 20 ms bins advanced in 10 ms time increments. For each event, at a given time point, we computed the posterior probability of all locations using the CA1 data and the left-arm and right-arm decoders, and combined them in a posterior probability matrix. As the decoders were not trained on time bins with zero spikes, the posterior probability for time bins with no spikes was set to a uniform distribution. We then used a line fitting algorithm similar to Davidson (Davidson et al., 2009) to determine the most likely linear trajectory from the posterior probabilities. That is, given a matrix of posterior probabilities of shape N (number of locations) by M (number of time bins) we first wrapped 10 locations on either side, then generated all possible combinations of start and end locations. To avoid trajectory events which represented only one location, we required trajectories to cover at least 2 bins and have a speed of at least 100 cm/s (which corresponds to requiring that the trajectory covers a distance of at least 2 cm in a time bin of 20 ms, roughly a fourth of the distance between two spatial bins). For each of the selected trajectories, we computed a ‘replay score’ (Davidson et al., 2009) by summing the posterior probabilities of 3 spatial bins around the trajectory line (the bin crossed by the line and one additional bin on either side, corresponding to roughly 25 cm) and expressed the score as a percentage of the total posterior probability. Finally, we selected the trajectory with the highest score. This was repeated for left-arm and right-arm decoders, and the fitted trajectory with the highest score was selected as the virtual trajectory and tested for statistical significance.

#### Statistical significance

We tested the significance of candidate events against three null distributions, a “column-cycle shuffle” (Davidson et al., 2009; Silva et al., 2015) which consists of applying a random circular shift to the probability estimates at each time point, a “unit identity shuffle”, which consists of randomizing the cell identity before computing the posterior probability matrix (Diba and Buzsáki, 2007; Foster and Wilson, 2006), and a “place field cycle shuffle” (Grosmark and Buzsáki, 2016; Ólafsdóttir et al., 2016), which consists of applying a random circular shift to the rate maps of the individual units before computing the posterior probability matrix. For a given candidate event, and for each shuffling method, we generated 300 surrogates, and assigned a score to each surrogate using the line fitting procedure to determine the best fit. We then computed a p-value as the fraction of surrogate replay scores which were higher than the observed score. Candidate events for which all three p-values were smaller than 0.05 were considered trajectory events. Even though the overlap between trajectory events and SWRs was incomplete (44 %), we consider it unlikely that many trajectory events took place during theta sequences. First, the movement speed of the animal’s head for trajectory analysis was low (< 3 cm/s, including sideways displacement) and second, visual inspection of the hippocampal LFP revealed only low or no theta rhythm during trajectory events. Nonetheless, theta sequences may also be generated during nearly immobile states (Gupta et al., 2012; Johnson and Redish, 2007), hence a strict association of the trajectory events presented here with SWR-associated replay is avoided here.

#### Distinction of forward and reverse trajectories

As in previous experiments (Bhattarai et al., 2020; Ólafsdóttir et al., 2016), the behavioral task imposed a specific running direction on the maze, thus making it impossible to determine directionality of trajectories based on rate maps obtained from bidirectional running (Ambrose et al., 2016; Davidson et al., 2009; Dragoi and Tonegawa, 2011; Wu and Foster, 2014). We determined whether virtual trajectories followed the running direction of the task (forward trajectory) or the opposite direction (reverse trajectory) based on the gradient of the fitted line (Bhattarai et al., 2020; Ólafsdóttir et al., 2016). During task performance, actual opposite direction runs were rare and firing patterns deviated considerably from those during forward runs (Bos et al., 2017); therefore, it is deemed unlikely that reverse trajectories, as reported here, predominantly reflect such opposite direction runs.

### Assessing coordination between hippocampus and perirhinal cortex

#### Data selection

For this analysis we selected recording sessions which had at least 10 trajectory events of the type considered and which contained at least one PER single unit. This resulted in 222 PER single units for all trajectory types combined (15 recording sessions, Fig. 4 and Supplementary Fig. 3 and 4). When forward and reverse events were considered separately, 189 units were identified (13 recording sessions, Fig. 4 and Supplementary 4). When considering only SWR-associated trajectories, we identified 201 units (14 sessions). When considering only trajectories not accompanied by an SWR, 177 units were listed (12 sessions, Supplementary Fig. 5).

#### Correlation between real and virtual rate maps

Using the method described by Ólafsdóttir (Ólafsdóttir et al., 2016) we constructed a real rate map, for each cell in PER, plotting spikes as a function of rat position during locomotion, and a virtual rate map, plotting spikes fired during hippocampal trajectory events to the reconstructed locations of the virtual trajectory. For the real rate map, we binned spikes in 200 ms non-overlapping bins. The firing-rate value of the map at a given location is the sum of all spikes fired in time bins assigned to the corresponding spatial bin divided by the total amount of time spent at that location. The same procedure was repeated for trajectory events (spikes binned in time windows of 20 ms bins advanced by 10 ms increments) to generate the virtual rate maps. Rate maps were smoothed with a Gaussian kernel with a standard deviation of 2 spatial bins. The virtual rate map was then compared to the real rate map using Pearson’s correlation. Firing rates plotted on maps were normalized to be between 0 and 1 for rendering purposes only (Supplementary Fig. 3).

#### Significance testing

To determine the statistical significance of the correlation between real and virtual rate maps, both at the population level and for individual cells, we constructed two null distributions (Ólafsdóttir et al., 2016). The first null distribution (‘rate map rotation’) was generated by shifting (and wrapping around at the edges) the real rate map to all possible spatial lags, and for each lag computing the correlation with the virtual rate map. The second one (‘spike vector rotation’) was obtained by shifting the vectors of temporally binned spikes in the trajectory event for each cell, where the amount of shifting was between 0 and the length of the spike vector minus one, and was chosen randomly and independently for each cell. A surrogate virtual rate map was then constructed from the shuffled event spikes, and correlated with the real rate map. This procedure was repeated 100 times for each neuron to obtain the null distribution of correlation values.

#### Debiased correlation coefficient (*r_d_*)

For every neuron, and for each of the null distributions (rate map rotation and spike vector rotation), we computed a debiased correlation score, as the observed correlation between the real and virtual rate maps minus the average of the correlation values of the surrogate distribution. For each null distribution, we computed a p-value as the fraction of surrogate correlations greater than or equal to the observed correlation. Single units for which both p-values were smaller than 0.05 were considered to be significantly coordinated with hippocampal trajectory events. We then assessed whether the amount of significantly coordinated units exceeded the number expected by chance (5% of the total) using a binomial test.

#### Temporal dynamics of cross-area coordination

To investigate the possibility of a temporal lag between representations in the HPC and PER, we applied a temporal shift to the perirhinal spikes during trajectory events, in increments of 10 ms, from -100 to 100 ms. For each increment, we pooled the *r*_*d*_ factors across cells and applied Wilcoxon signed-rank test. We then corrected for multiple comparisons using the Benjamini-Hochberg method, and repeated the procedure for both null distributions (rate map rotation and spike vector rotation). Lastly, we required the distribution of *r*_*d*_ at a given temporal shift to be significant with respect to both null distributions.

### Statistical analysis

The details of the statistical comparisons, the definition of center and dispersion are contained in the figure legends. Throughout the text we report our findings as median and interquartile ranges unless stated otherwise. Data selection criteria for position decoding, trajectory event detection, and cross-area coordination are listed at the beginning of each section. In general, we applied non-parametric statistical tests: Wilcoxon signed-rank test (paired data) and Mann-Whitney’s U test (unpaired data). We generated confidence intervals using bootstraps. Whenever multiple comparisons were performed, we adjusted p-values using Bonferroni’s correction prior to reporting and assessing significance, and with the Benjamini-Hochberg procedure in the temporal analysis of cross-area coordination (Figs. 4E-F and Supplementary Fig. 4 and 5). The cross-validation strategy for position decoding and the permutation methods to assess the significance of trajectory event detection and coordination are detailed in the corresponding sections. For the comparison of the distributions of start and end points of trajectory events across task phases (Fig. 3F), we subdivided the maze in 8 macro bins, in order to guarantee enough observations in each bin for the Chi-square test (the macro bins were used for the statistical test only, not for plotting). For the computation of confidence intervals we relied on the Seaborn library (https://github.com/mwaskom/seaborn).

## Results

The animals performed a visual discrimination task on a figure-8 maze (Fig. 1A-B). Each trial began with the animal situated in the central arm of the maze. Following an auditory cue that signaled trial onset, two images, a target and a distractor stimulus, appeared on the two screens positioned in front of the central arm. Rats had to travel along the arm of the maze corresponding to the side of the target stimulus to earn a reward (Bos et al., 2017). Each trial was subdivided into three task phases. The ‘ITI’ (inter-trial interval) was defined as the time after the animal had left the reward site in the previous trial and before the stimulus presentation of the upcoming trial. The ‘Image’ phase consists of the time between the visual appearance of the stimuli and the removal of the front barrier (which allowed the animals to start moving towards the arms). Lastly, the ‘Reward’ phase began when the animal crossed a photobeam placed at the reward site (and, in the case of correct trials, received food pellets), and ended as they left the reward site to return to the central arm. Overall behavioral performance was above chance level (59.2 ± 0.5% correct trials, mean ± s.e.m.; p<0.001, Wilcoxon signed-rank test). Neuronal ensembles were recorded simultaneously from hippocampal area CA1 (N=557 cells) and PER (N=238 cells; see Materials and Methods).

### Decoding of the animal’s location and global positioning in maze arms

We first examined whether the actual global and local positioning of the animal on the maze could be reliably reconstructed from the joint spiking activity of hippocampal or perirhinal ensembles using a Bayesian decoder (Davidson et al., 2009; Zhang et al., 1998). This provides the basis for our subsequent analysis of different types of CA1 trajectory events throughout the task, and of whether perirhinal representations cohere with the trajectories represented by area CA1 ensembles. CA1 ensembles allowed for precise reconstruction of the animal’s location (Fig. 1C), and the decoding of location was significantly better than chance in all recordings (p<0.001). Perirhinal cells recorded in this task were previously found to display broad, sustained firing fields for specific subsections of the maze (Bos et al., 2017). Due to these characteristic broad firing fields, fine-grained prediction of location was not successful. Instead, the Bayesian decoder was trained to predict whether at a given point in time the animal was situated on the left, central, or right arm of the maze. PER firing patterns allowed a reliable decoding of maze arm (Supplementary Fig. 1B), with balanced accuracy score of 0.46, [0.46 - 0.48] (median and interquartile range), and this decoding was significantly better than chance in all 11 sessions considered (p<0.001). Thus, while hippocampal ensembles predicted the precise location of the animal, PER firing patterns could be exploited by the decoder to determine the global maze segment the animal was located on.

### Detection of virtual trajectories by CA1 ensembles

Next, we investigated the neural coding of virtual trajectories by hippocampal ensembles during the full task period, including the ITI, by identifying population burst events (PBEs) as periods in which the animal was relatively immobile (head movement speed (< 3 cm/s, Fig. 1D, see Materials and Methods) and the intensity of multi-unit activity (MUA) of putative CA1 pyramidal cells exceeded 3 s.d. above the mean. We employed a Bayesian decoding algorithm (Davidson et al., 2009; Zhang et al., 1998) together with a line-fitting procedure to identify virtual trajectories (Fig. 2). We trained two Bayesian decoders, using rate maps calculated for all left and right trials, respectively. The spikes of a candidate PBE were first divided in 20 ms bins with a sliding window of 10 ms (left panels with grey bins in Fig. 2), then for each time bin a Bayesian decoder assigned a probability to each spatial bin, resulting in a posterior decoding probability matrix (right panels with blue bins in Fig. 2). For each decoder, the trajectory line was identified as the line along which posterior probability was maximized. The line with the highest overall posterior probability was then tested for significance (p<0.05) against three shuffled distributions (see Materials and Methods). Of 4633 putative events detected across 16 recording sessions, 1261 (27.2%) were found to be significant against all three shuffled distributions and considered (virtual) trajectory events. The slope of the fitted line was used to distinguish between a forward trajectory, when the decoded path followed the direction of running during the task, and a reverse trajectory, when it ran in the opposite direction. Hippocampal replay and theta sequences typically involve time-compressed representations (Gupta et al., 2012; Nádasdy et al., 1999). We estimated the compression factor as the speed of the trajectory divided by the average running speed of the animal on the maze arms in the corresponding session, and we found an average compression factor of 5.8 (across all task phases together, median and interquartile range: 3.8 [2.5, 7.1]). This is of the same order of magnitude as previously reported for CA1 during sleep (Ji and Wilson, 2007; Lansink et al., 2009; Lee and Wilson, 2002; Nádasdy et al., 1999) or during theta sequences in the awake state (Dragoi and Buzsáki, 2006; Maurer et al., 2012; Pezzulo et al., 2017).

**Figure 2.**
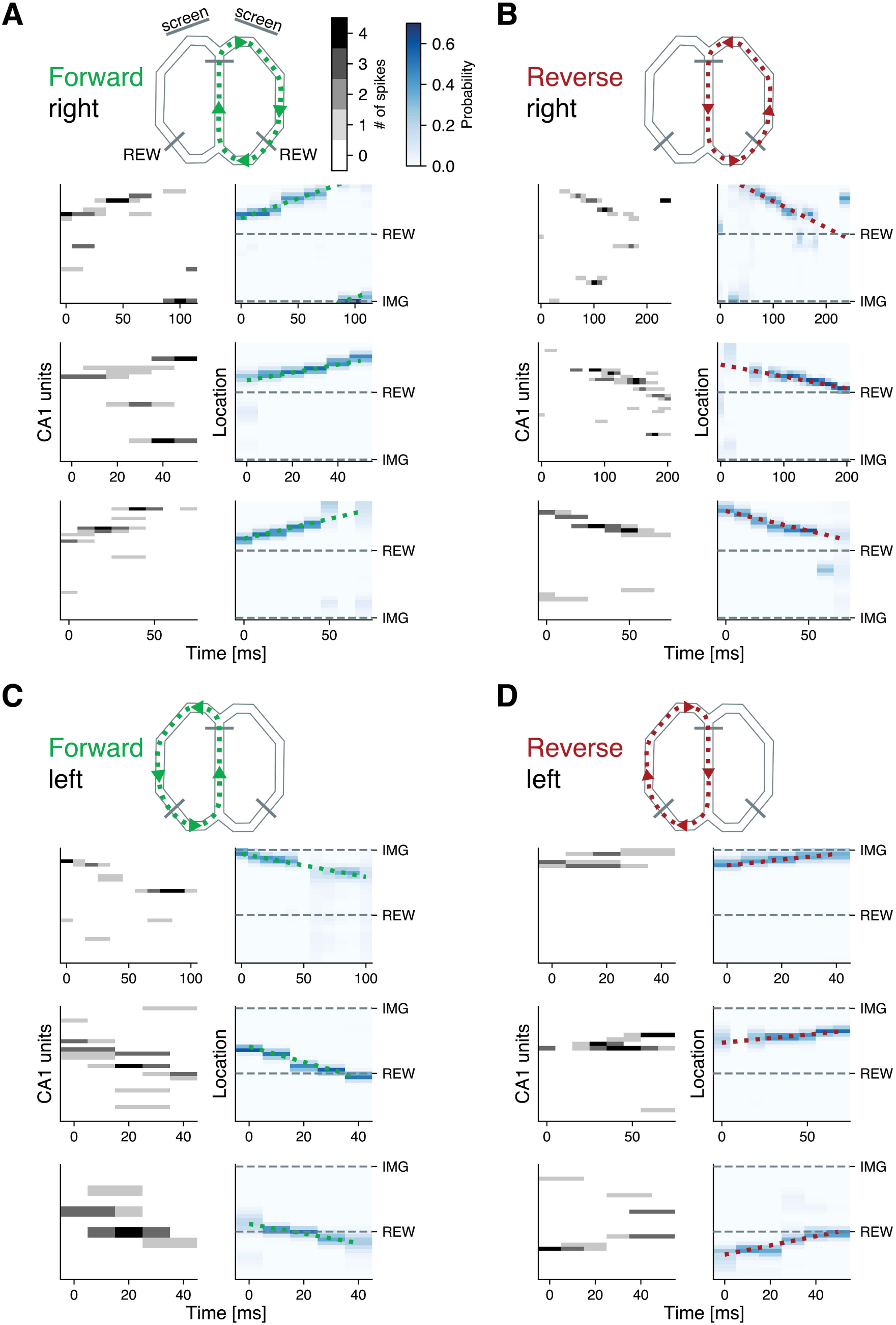
Examples of forward and reverse hippocampal place cell sequences in the awake state. **(A-D)** Examples of virtual trajectory events detected during task performance and intertrial intervals. Left panels, grayscale: binned spikes of putative pyramidal CA1 neurons sorted by location of peak firing. Right panels, blue heatmap: posterior probability matrix of animal location generated via Bayesian decoding; green and red dotted lines indicate the fitted virtual trajectory. Sequences reiterated non-local trajectories on either the right or left maze arm, and the virtual trajectory could either follow the direction of task running (‘forward’ sequence) or the opposite direction (‘reverse’ sequence): (A) forward sequences of right-arm trajectories, (B) reverse sequences of right-arm trajectories, (C) forward sequences of left-arm trajectories and (D) reverse sequences of left-arm trajectories. Example sequences were not selected based on task phase. More examples in Supplementary Fig. 2.

### Organization of hippocampal trajectory events across task phases

We next analyzed how virtual trajectories generated by CA1 neurons were distributed across the maze and relative to the animal’s position during different task phases. Even though this analysis only addresses hippocampal sequences, regardless of PER activity, we consider it important to present because CA1 trajectories have been rarely characterized across distinct phases of a stimulus-driven and spatial task including an ITI period. We hypothesized that visual image presentations would trigger the generation of future path options and thus would enhance the frequency of trajectory events, while additional events may occur during ITIs or the reward phase. We first computed a trajectory rate as the number of CA1 trajectories per minute of relative immobility, and found that the ITI had a significantly higher rate (4.9 events per minute on average) than the Image and Reward phases (2.2 and 2.0 events per minute on average, respectively, Fig. 3A; p<0.05, Wilcoxon signed-rank test). This indicates that virtual trajectories were more frequent during the rest phase than during epochs of task engagement, including the image presentation phase. Similar trajectory rates were found in correct and incorrect trials (data not shown).

**Figure 3.**
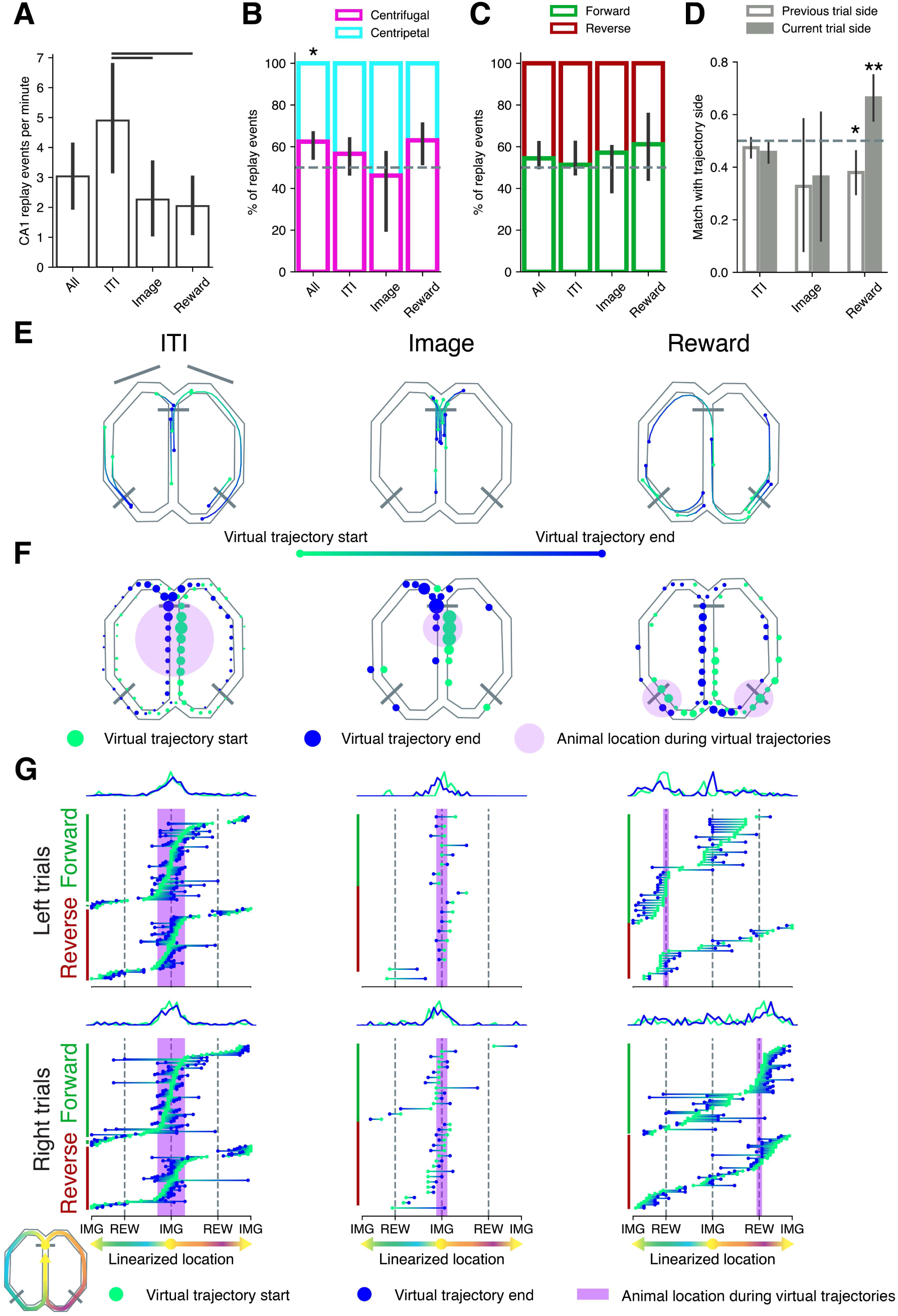
Characteristics of hippocampal trajectory events are heterogeneous across behavioral phases. (A) Trajectory rate (number of events per minute of relative immobility) for the different task phases, averaged across sessions. Error bars indicate 95% bootstrapped confidence intervals (CIs). Horizontal lines indicate significance at p<0.05 (Wilcoxon signed-rank test). ITI: inter-trial interval; Image: stimulus presentation phase; Reward: food consumption and/or lingering at reward site. **(B)** Percentage of centrifugal trajectories (moving away from the animal’s actual position during the virtual trajectory, fuchsia) and centripetal trajectories (moving towards the animal, light blue) per task phase (median across sessions with at least 5 trajectory events for each given phase, error bars indicate 95% bootstrapped confidence intervals). * indicates p<0.05, one-sample Wilcoxon signed-rank test. **(C)** As (B), but for the percentage of forward (green) and reverse (red) events. Differences were not significant. **(D)** Accordance between the side of the maze chosen by the animal in the current (filled grey bars) and previous (white bars) trial and the side on which virtual trajectories terminated, for the different task phases. Match was computed using a balanced accuracy score; error bars indicate 95% bootstrapped CIs, * indicates p<0.01, ** indicates p<0.001. A match value below 0.5 indicates that CA1 ensembles avoid representations of trajectories on the side of the maze visited in the previous trial. **(E)** Examples of 8 randomly chosen virtual trajectories per task phase. Throughout the figure, light green indicates start of virtual trajectory, blue indicates end. **(F)** Distribution of start and end locations of virtual trajectories for each task phase. Start and end locations have been plotted at different sides of each maze arm for rendering purposes only. Dot diameter indicates proportion of trajectory events which start or end at a given location. Distributions of start and end locations were different across task phases (chi-square test, p<0.001 for all comparisons). **(G)** Trajectories on a linearized axis, sorted by location and trajectory direction, and split based on maze arm chosen by the animal; for ITI and Image phases this corresponds to upcoming choice). Purple bands indicate approximate animal position at the time of the trajectory event. Top panels: distributions of start and end locations of the trajectories (green and blue lines, respectively).

Next, we considered the spatial direction of virtual trajectories relative to the actual location of the animal. We labelled trajectories as ‘centrifugal’ (moving away from the animal) when the start point of the trajectory was closer to the actual position of the animal at the time of the event than the end point of the same trajectory, and as ‘centripetal’ (moving towards the animal) when the opposite was true. We found a significant predominance of trajectories moving away from the animal (Fig. 3B, percentage of centrifugal events in the full task, 62% [55% - 66%]; p<0.01, Wilcoxon signed-rank test). This effect was also reflected in a trend that was found in the ITI and Reward trajectories when considered separately (57% [48% - 64%] and 63% [53% - 70%]), while centrifugal events during the Image phase tended to be under-represented (46%, [33% - 52%]). Functionally, there is growing support for a distinct role of reverse and forward awake replay: reverse replay has been linked to memory consolidation and retrospective evaluation (Ambrose et al., 2016; Foster and Wilson, 2006; Michon et al., 2019; Shin et al., 2019), while forward replay may enable exploration of future outcomes (Pfeiffer and Foster, 2013; Shin et al., 2019; Wu et al., 2017). Thus, forward and reverse trajectories may be utilized selectively depending on task requirements. Across our full task, we found a balanced repertoire of forward and reverse trajectories (Fig. 3C, percentage of forward events: 54% [50% - 62%], p=0.4, Wilcoxon signed-rank test), with a slight but non-significant prevalence of forward events. This was also the case when considering individual task phases separately. We also contrasted forward and reverse trajectories using other metrics, including replay scores (see Materials and Methods), percentage of recruited neurons, event duration, maximum jump distance, sharpness (Silva et al., 2015) and compression factors, but found no salient differences.

Hippocampal sequences have been proposed as a route planning mechanism, as they can reflect future behavioral paths (Pfeiffer and Foster, 2013) or avoided paths (Wu et al., 2017). We explored the relationship between the arm on which CA1 trajectories ended during the different behavioral phases and the path chosen by the animal in the current or previous trial. During the Image phase, when the animals were exposed to a visual stimulus, the end point of the virtual trajectory generally failed to predict the maze arm corresponding to the animal’s subsequent choice; similarly, trajectories during the ITI did not over-represent the arm of the maze that the animal had arrived from (Fig. 3D). These results did not change when considering forward and reverse trajectories separately. In contrast, during the Reward phase, trajectories were more likely to end on the arm that the animal was currently on (66% [57% - 75%], p<0.001), and more likely to avoid the trajectory side chosen in the previous trial (38% [29% – 46%], p<0.01).

Next, we analyzed the fine-grained spatial content of virtual trajectories. Examples of trajectories for the ITI, Image and Reward phases (Fig. 3E) suggest that during subsequent phases different portions of the maze were represented by hippocampal sequences. During the ITI and Image phases, start and end points of the virtual trajectories were predominantly situated in the middle lane, with a higher concentration of end points around the bifurcation of the two arms (Fig. 3F). Trajectories during Reward were often positioned closer to the reward sites and in the middle lane. The distributions of start and end locations over the maze (Fig. 3F) differed significantly across task phases (p<0.001 for all comparisons, Chi-square test).

We further examined the spatial organization of sequence representations by plotting trajectories on a linearized axis, grouped by trajectory direction (forward vs. reverse) and side of the trial (left vs. right, for ITI and Image phases we considered the upcoming chosen side), together with densities of the start and end locations of the trajectories (Fig. 3G, upper row). During ITI and Image, the animal was positioned in the middle lane, and most of the start and end points of the trajectories occurred in the vicinity of the animal, but trajectories during the Image period appeared more concentrated around the animal’s location. Thus, the hypothesis about image onsets triggering enhanced occurrence of place cell sequences is not supported by our data. Trajectories in the reward phase were widely distributed across the maze, with increased density of start and end points near the reward locations, but also a notable subset starting at the Image (IMG) location and running forward into the arms.

### Independence of hippocampal place cell sequences from perirhinal activity

We next investigated whether PER activity co-varied with hippocampal place cell sequences. We first asked whether the onset of virtual trajectories was accompanied by changes in PER firing. To compute the firing-rate modulation of individual PER neurons by CA1 trajectory events, we identified a baseline as the interval from -600 to -100 ms relative to trajectory onset. The maximum firing rate during each trajectory event, z-scored with respect to the baseline, was then taken as the modulation. Neurons were considered significantly responsive if the average response across trajectory events exceeded 2 standard deviations. Overall, only very small fractions of PER neurons appeared to modulate their firing during trajectory events (0% when considering all trajectory events, 1% when considering forward trajectory events, 3.6% when considering reverse trajectory events). Mean z-scored firing rates (averaged across all PER neurons) also showed no clear timelocked modulations (Fig. 4A-B).

**Figure 4.**
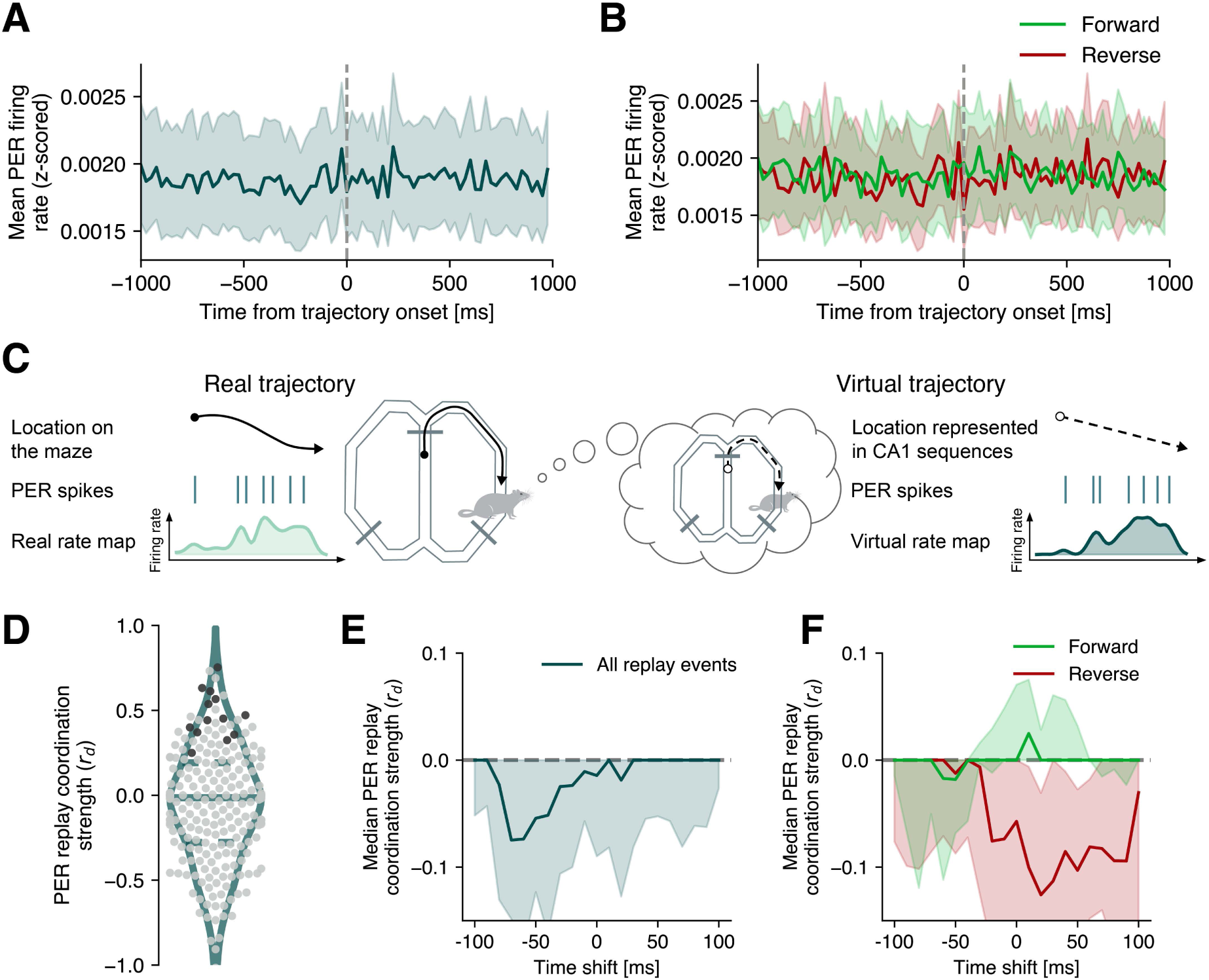
Perirhinal activity is not engaged by hippocampal trajectory events. (A) Firing rate (z-scored) averaged across all PER neurons aligned to the onset of hippocampal place cell sequences. **(B)** Same as (A) but computed separately for forward and reverse trajectories. **(C)** Schematic of the virtual rate map method used to assess coordinated PER-HPC activity. For a given PER neuron, we computed the real rate map using the animal’s linearized position (across the whole session). We then computed the virtual rate map with the spikes the cell fired during hippocampal trajectories, using as positions the trajectory points represented by CA1. A high correlation between the two maps indicates that the cell’s firing pattern is consistent with CA1 during trajectory events. **(D)** Replay coordination strength measured with the debiased correlation coefficient (*r*_*d*_) for individual PER units. Solid and dashed horizontal lines within violin plots indicate median and quartiles of distributions. Dark dots correspond to significantly correlated units (considering both permutation methods). Here and throughout the rest of the figure, debiasing is performed with spike vector rotation. For results using the rate map rotation method, see Supplementary Fig. 4. **(E)** PER-HPC coordination strength (median *r*_*d*_) as a function of time shift relative to CA1 spikes. Shaded bands indicate 95% CIs. **(F)** Same as (E), computed for forward and reverse events separately.

Lack of a marked increase or decrease in firing rates, however, does not rule the possibility of replay coordination, especially if PER is involved in a content-based recapitulation. To address this possibility, for every PER unit, we built a ‘real’ rate map plotting its spikes as a function of the animal’s actual position during locomotion, and a ‘virtual’ rate map (Ólafsdóttir et al., 2016), plotting perirhinal spikes fired during hippocampal trajectories as a function of the location within the non-local trajectory decoded from CA1 ensembles (Fig. 4C). A strong resemblance between the real and virtual rate maps indicates that, during the trajectory event, the representation coded by a perirhinal cell (be it spatial or otherwise) is consistent with the spatial representation of the HPC, providing evidence for coordinated activity. We identified neurons which had a significant correlation between real and virtual rate maps (‘coordinated’ neurons) using two permutation procedures (‘spike vector rotation’ and ‘rate map rotation’, see Materials and Methods). Although some individual units showed a high correlation between real and virtual rate maps (Supplementary Fig. 3), only a small proportion were identified as significant by our method (Fig. 4D, 13 out of 222 neurons, 5.9%, p=0.5, binomial test). As a measure of replay coordination, we defined a debiased correlation coefficient (*r*_*d*_) as the observed correlation between the real and virtual rate map of a given unit minus the average correlation of the surrogate data. This coefficient was computed separately for the two permutation methods: in Figure 4 and in the remainder of this section we discuss results pertaining to the spike vector rotation (for analogous results with rate map rotation, see Supplementary Fig. 4). Following Ólafsdóttir et al. (2016), we also computed a percentile rank of the correlation given each of the null distributions, and used Wilcoxon signedrank test to determine if the percentile ranks of the population were significantly different from the expected median rank of 50%. For PER cells, we found a median rank of 65.5 [31.5 - 87.9], with p<10^-4^ for rate map rotation, and of 53.0 [0.0-94.5] with p=0.345 for spike vector rotation). Thus, significance was reached for only one of the two null distributions.

### Temporal relationship of perirhinal activity relative to hippocampal trajectories

We next considered the possibility of a temporal lag between hippocampal sequences and perirhinal activity: memory operations may be initiated by the HPC before reaching other structures (Ji and Wilson, 2007; Lansink et al., 2009; Ólafsdóttir et al., 2016), but can also originate in the cortex (O’Neill et al., 2017; Rothschild et al., 2017). We applied a time shift to the neocortical spikes of all cells in each trajectory event, from -100 to 100 ms in steps of 10 ms. For every time shift, we recomputed the virtual rate maps using the shifted spikes, and followed the procedure outlined above to compute the *r*_*d*_ of all neurons, for both permutation methods. Correlation at a given time point was considered significant if both distributions were different from an expected median of zero.

We found that the absence of a significant perirhinal coordination at t=0 extended to all other lags with respect to CA1 sequences (Fig. 4E), also when considering forward and reverse trajectories separately (Fig. 4F). Thus, while perirhinal representations correlate with the animal’s location (and thus with CA1 place cell activity) during locomotion (Supplementary Fig. 1), this correspondence is not preserved when virtual trajectories are sampled during CA1 place cell sequences. We also asked whether perirhinal coordination could be restricted to CA1 trajectory events accompanied by SWRs, but found no evidence of correlation even when considering only SWR-associated trajectories (Supplementary Fig. 5).

Perirhinal reactivation might be recruited selectively to recapitulate important episodic memories such as those associated with reward. We thus computed replay coordination in the subset of virtual trajectories which occurred during the Reward phase of correct trials, or in the ITI following correct trials, but again found no significant coordination (data not shown). Thus, while perirhinal activity is correlated to the animal’s current location, allowing successful decoding of the arm of the maze currently occupied, these representations are not evoked (or only very marginally) in coordination with hippocampal sequences.

## Discussion

We found that CA1 sequences comprised a rich variety of events, characterized as forward versus reverse, local versus remote and centrifugal versus centripetal sequences. Overall, centrifugal sequences were prevalent, and a balanced mixture of forward and reverse sequences across task phases was found. During stimulus presentation (Image phase), virtual trajectories sampled both the maze arm that animals were about to choose as well as the non-chosen side. At reward sites, CA1 sequences over-represented the arm of the maze that the animal was on, and selectively under-represented the arm of the maze visited in the previous trial. PER cells did not display marked firing rate modulations around the onset of CA1 sequences, and only a small fraction showed activity which was coordinated with CA1 trajectories.

The implications of the broad repertoire of CA1 trajectory types we report for the overall task need to be considered. Under certain behavioral conditions, forward CA1 replay sequences correlate with the path the animal will take (Ito et al., 2015; Pfeiffer and Foster, 2013; Wood et al., 2000). In general, however, they do not strictly predict upcoming behavior (Carey et al., 2019; Gillespie et al., 2021; Wu et al., 2017), excluding a one-to-one relationship between forward sequences and future paths. The heterogeneous CA1 trajectories we detected during ITI and Image epochs are consistent with the absence of a simple predictive relationship between a virtual trajectory and the recently travelled path or upcoming behavioral choice (Pfeiffer and Foster, 2013; Wu et al., 2017). Indeed, viewing a particular image at the start of each trial was not coupled to sequence events representing the path that the animal would choose next (Fig. 3D). Moreover, task-relevant images did not appear to enhance the rate of CA1 trajectory events (Fig. 3A), hence image viewing does not trigger an unusual generation of replays. On the other hand, trajectory events during Reward epochs were biased to the maze arm the animal occupied (Fig. 3D), and they especially covered the portion of the maze between the reward well and the central arm, whereas paths leading to the reward location along the maze were less represented.

Previous research has linked hippocampal replay with reward, showing how reward increases replay activity (Singer and Frank, 2009) and how this modulation mostly boosts reverse replay (Ambrose et al., 2016; Michon et al., 2019). However, the relationship between reinforcement and (reverse) CA1 replay is not straightforward: both forward and reverse replay can be biased towards feared locations (Wu et al., 2017) and non-preferred outcomes (Carey et al., 2019). Preplay of reward-related sequences was found to be both forward and reverse (Ólafsdóttir et al., 2015), and reward was shown to modulate forward and reverse replays differently (Bhattarai et al., 2020). Our finding that in the Reward phase both forward and reverse trajectories appeared in similar amounts and were biased towards the currently occupied reward location are thus compatible with previous findings, including those which characterized how reward dynamically structures hippocampal replay depending on task requirements (Sosa and Giocomo, 2021).

In addition to the HPC, replay and related virtual trajectories have been shown in a number of different brain structures: visual cortex (Ji and Wilson, 2007), auditory cortex (Rothschild et al., 2017), posterior parietal cortex (Qin et al., 1997), ventral striatum (Lansink et al., 2009; Pennartz et al., 2004), deep (Ólafsdóttir et al., 2016; Ólafsdóttir et al., 2017) and superficial layers of MEC (O’Neill et al., 2017), basolateral amygdala (Girardeau et al., 2017), ventral tegmental area (Gomperts et al., 2015), cingulate cortex (Remondes and Wilson, 2015) and medial prefrontal and orbitofrontal cortex (Kaefer et al., 2020; Rusu et al., 2024; Shin et al., 2019; Tang et al., 2017). Cross-structural replay is not a general phenomenon; rather, the involvement of brain areas other than the HPC appears to be selective to awake, centrifugal replay for VTA (Gomperts et al., 2015), to replay trajectories actualized by the animal for medial PFC (Shin et al., 2019), to forward replay in sleep (Ólafsdóttir et al., 2016) and extended (‘disengaged’) immobility periods during wakefulness (Ólafsdóttir et al., 2017) for MEC deep layers. In our current study, the lack of evidence for coordination between hippocampal trajectories and PER activity corroborates the idea that the involvement of other structures (even nearby parahippocampal structures) is not a general feature of hippocampal memory operations. If this coordination occurs, it appears to be subject to modulation by the specific area, the type of hippocampal trajectory and the requirements of the task at hand. Furthermore, reactivation of episodic or spatial memories may not be initiated solely by the hippocampus: superficial layers of the medial entorhinal cortex (MEC) display reactivation sequences which occur independently of the hippocampus, possibly acting as a parallel system and recruiting a different coalition of brain structures (O’Neill et al., 2017), while input from MEC layer III contributes to extended hippocampal SWRs in the quiet awake state (Yamamoto and Tonegawa, 2017). The characteristics of MEC replay and its relationship with hippocampal replay, as well as the role of the hippocampus in shaping or coordinating cortical reactivations (Aly et al., 2022) are not well known, but we may speculate that PER is involved in memory operations without a simple direct correspondence with hippocampal sequences. For instance, the activity state of the PER may be strongly modulated by novelty or surprise in trial outcome (Fiorilli et al., 2024). In our task, the finding that CA1 can perform memory operations in the form of place cell sequences independently of PER is also in line with recent work on this dataset which showed PER spiking activity to be weakly coupled to HPC and visual and somatosensory cortical areas (Dorman et al., 2023).

## Acknowledgements

We would like to thank Julien Fiorilli for his comments on the manuscript. We acknowledge the software tools provided by Kenneth Harris (University College London; Klustakwik), A. David Redish (University of Minnesota, Minneapolis; MClust), Michael Waskom and the Seaborn development team. The data is available on the EBRAINS platform (Bos, 2019), DOI: 10.25493/1PEP-K9F.

## Author contributions

Conceptualization, C.P. and P.M.

Investigation, J.B. and M.V.

Formal analysis, P.M., J.B., M.V.

Methodology, P.M., J.B., M.V., C.P.

Software, P.M.

Supervision, C.P.

Funding Acquisition, C.P.

Writing – original draft, P.M. and C.P.

Writing – review and editing, P.M. and C.P.

---

**Supplementary Figure 1:**
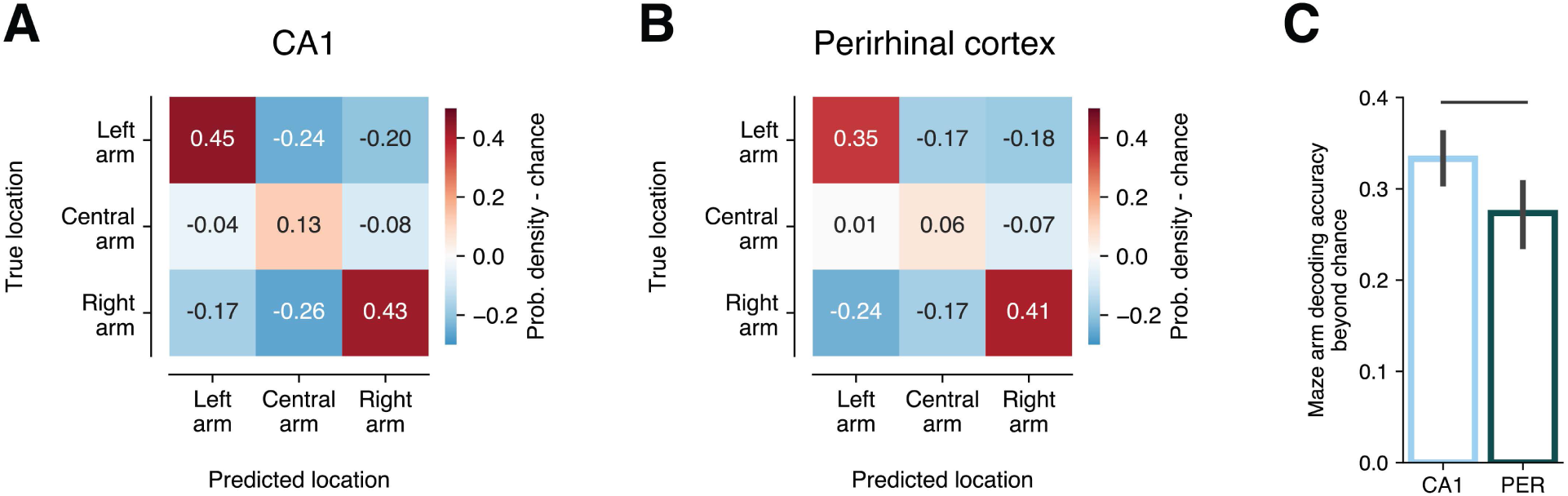
Decoding of the animal’s actual location and global positioning in maze arms. (A-B) Confusion matrices for the prediction of animal location grouped in segments of the maze environment from CA1 (A) and PER (B) ensembles, averaged across sessions (N=17 and N=11 for CA1 and PER, respectively). High positive values along the diagonal from top left to bottom right indicate a decoding performance beyond chance level. Decoders were typically less able to correctly identify the central arm, probably because firing patterns generated on this arm were less separable from patterns occurring on the left and right arms (cf. Bos et al., 2017). Decoding from PER ensembles was significantly better than chance in all 11 recording sessions considered. **(C)** Balanced accuracy scores for maze arm decoding for CA1 and PER (averaged across sessions). Error bars indicate 95% CIs. Horizontal bar indicates p<0.05 (Mann-Whitney U-test).

**Supplementary Figure 2.**
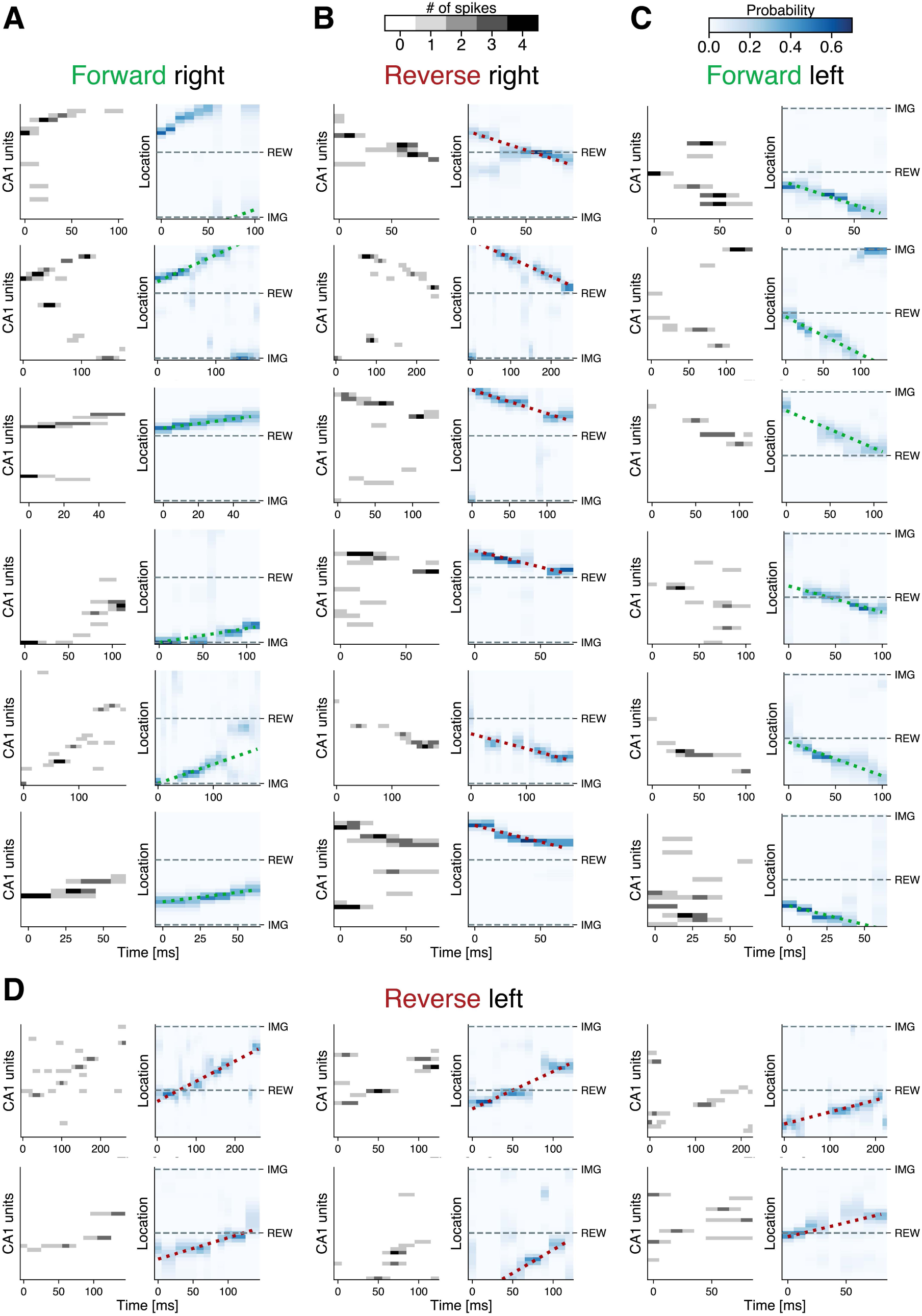
Additional examples of forward and reverse hippocampal place cell sequences in the awake state. (A-D) Examples of CA1 sequences (as in Fig. 2). Left panels, grayscale: binned spikes of CA1 neurons sorted by location of peak firing. Right panels, blue heatmap: probability matrix of animal location generated via Bayesian decoding; green and red dotted lines: fitted virtual trajectory. (A) Forward sequence of right-arm trajectories, (B) reverse sequence of right-arm trajectories, (C) forward sequence of left-arm trajectories and (D) reverse sequence of left-arm trajectories.

**Supplementary Figure 3:**
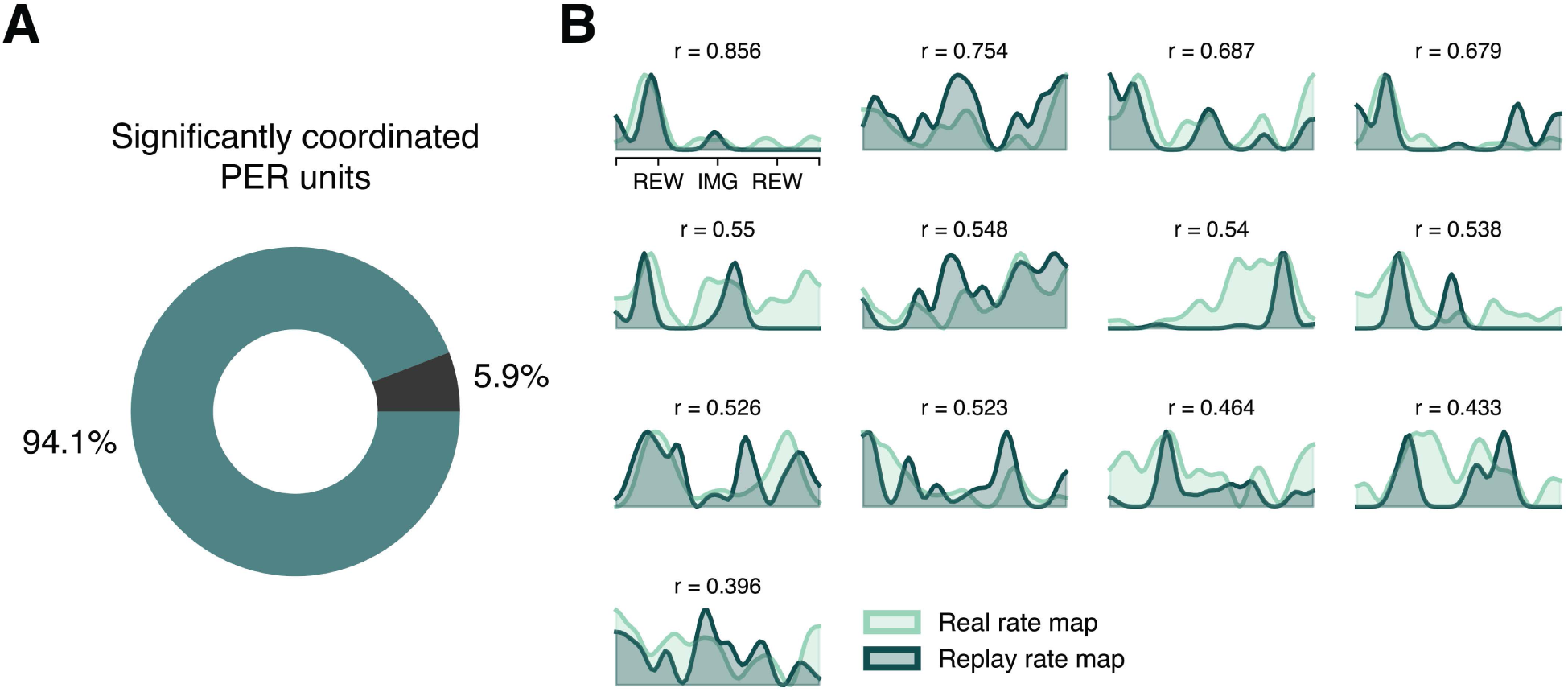
Activity of a small fraction of perirhinal units is coordinated with hippocampal virtual trajectories. (A) Percentage of coordinated perirhinal units (13 out of 222, 5.9%, p=0.5, binomial test). Coordination with CA1 trajectories was determined with two permutation methods, rate map rotation and spike vector rotation, and by requiring a cell to be significant against both null distributions. Despite the significance found for 5,8 % of the PER population, this percentage did not exceed the percentage of cells expected to show significance by chance. **(B)** Real and virtual rate maps for the 13 PER units showing coordination with CA1 trajectories. Values are Pearson correlation coefficients (r).

**Supplementary Figure 4:**
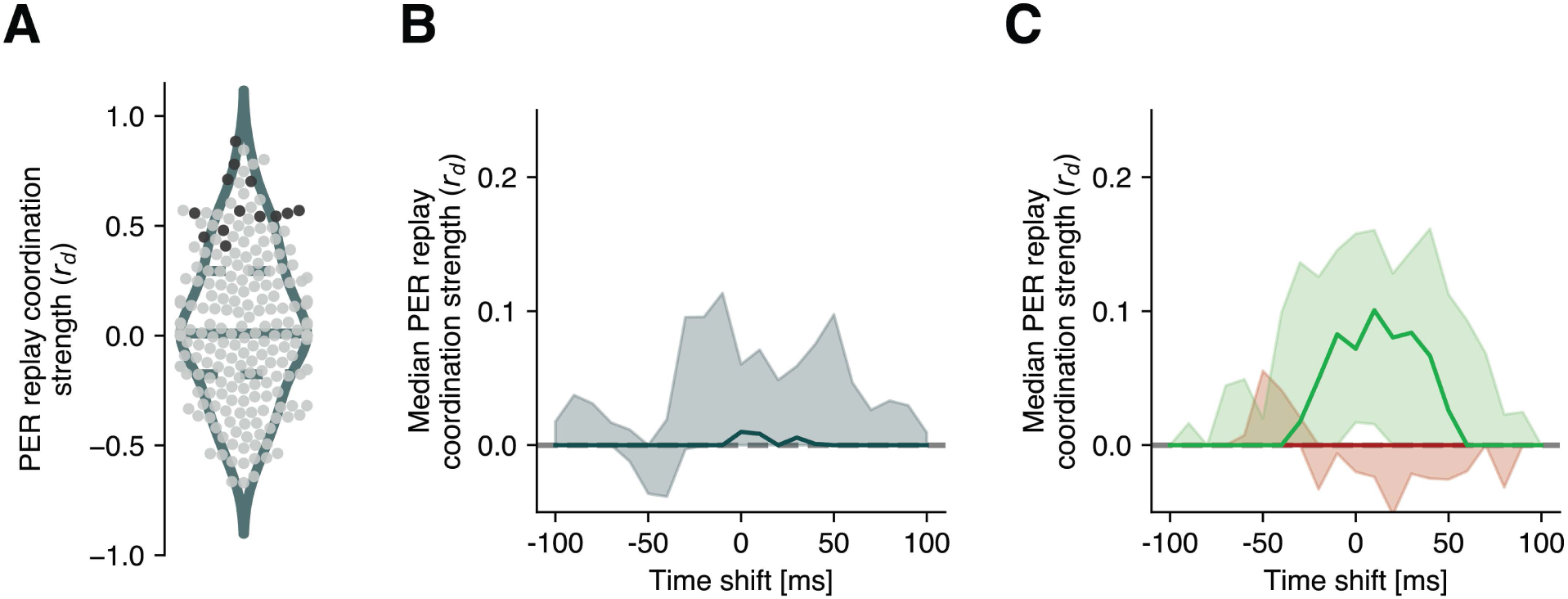
Perirhinal-CA1 coordination using the null distribution derived from rate map rotation. (A) Replay coordination strength measured with the debiased correlation coefficient (*r*_*d*_) for individual PER units. Solid and dashed horizontal lines within violin plots indicate median and quartiles of distributions. Dark dots correspond to significantly correlated units (considering both permutation methods). Debiasing (here and throughout the figure) is performed with rate map rotation. (B) PER-HPC coordination strength (median *r*_*d*_) as a function of time shift of the PER firing pattern relative to CA1 spikes. Shaded bands indicate 95% CIs. No significant deviations from r_d_=0 were found. (C) Same as (B), computed for forward and reverse events separately. No significant deviations from r_d_=0 were found.

**Supplementary Figure 5:**
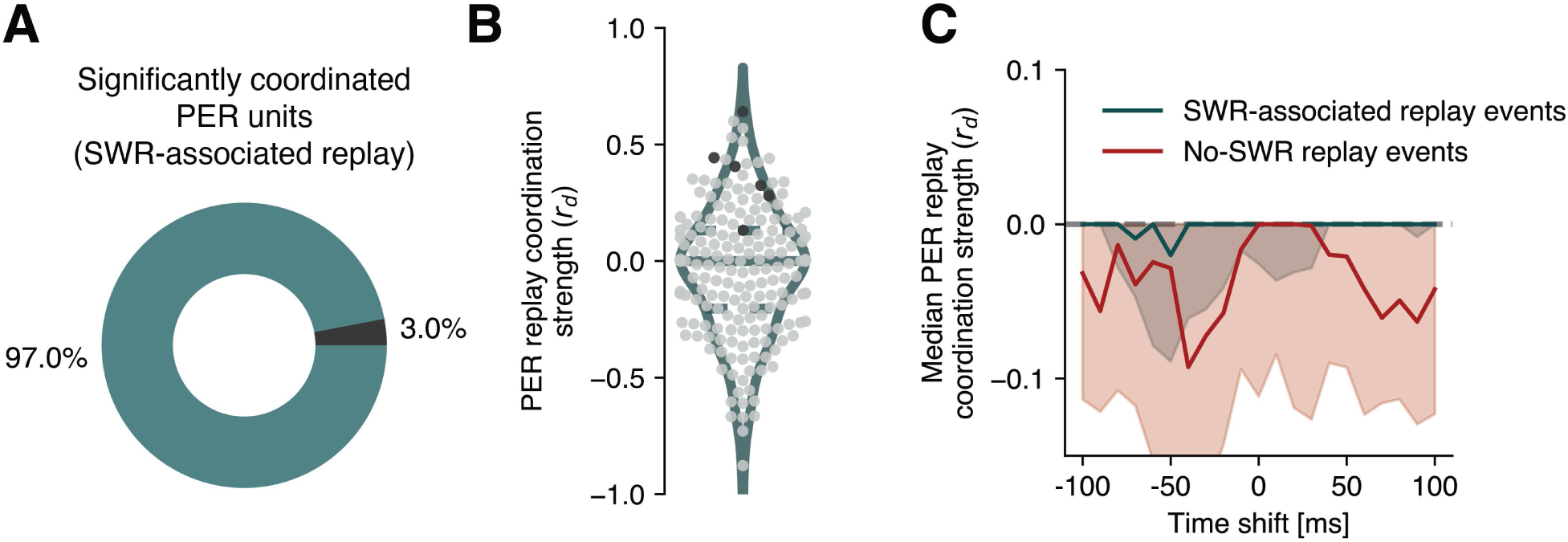
Perirhinal-CA1 replay coordination in sequences associated with sharp wave ripples. (A) Percentage of coordinated perirhinal units (6 out of 201, 3.0%, p=0.3, binomial test) computed on the subset of virtual CA1 trajectories co-occurring with sharp wave ripples (SWRs). Coordination with CA1 trajectories was determined with two permutation methods, rate map rotation and spike vector rotation, and by requiring a cell to be significant against both null distributions. **(B)** Replay coordination strength measured with the debiased correlation coefficient (*r*_*d*_) for individual PER units, computed on SWR-associated sequences. Solid and dashed horizontal lines within violin plots indicate median and quartiles of distributions. Dark dots correspond to significantly correlated units (considering both permutation methods). Throughout this figure, debiasing is performed with spike vector rotation, although debiasing with rate map rotation led to analogous results (not shown). **(C)** PER-HPC coordination strength (median *r*_*d*_) as a function of time shift relative to CA1 spikes, computed separately on virtual trajectories which were (or were not) accompanied by an SWR. Shaded bands indicate 95% CIs. Debiasing with rate map rotation led to analogous results (not shown).

